# Addressing preferred orientation in single-particle cryo-EM through AI-generated auxiliary particles

**DOI:** 10.1101/2023.09.26.559492

**Authors:** Hui Zhang, Dihan Zheng, Qiurong Wu, Nieng Yan, Zuoqiang Shi, Mingxu Hu, Chenglong Bao

**Affiliations:** Qiuzhen College, Tsinghua University, Beijing, China.; Yau Mathematical Sciences Center, Tsinghua University, Beijing, China.; Yanqi Lake Beijing Institute of Mathematical Sciences and Applications, Beijing, China.; State Key Laboratory of Membrane Biology, School of Life Sciences, Tsinghua University, Beijing, China.; Shenzhen Medical Academy of Research and Translation (SMART), Shenzhen, China.; Beijing Frontier Research Center for Biological Structure (Tsinghua University), Beijing, China.; Tsinghua-Peking Joint Center for Life Sciences, Tsinghua University, Beijing, China.; School of Life Sciences, Tsinghua University, Beijing, China.; Department of Molecular Biology, Princeton University, Princeton, NJ, USA.

## Abstract

The single-particle cryo-EM field faces the persistent challenge of preferred orientation, lacking general computational solutions. We introduce cryoPROS, an AI-based approach designed to address the above issue. By generating the auxiliary particles with a conditional deep generative model, cryoPROS addresses the intrinsic bias in orientation estimation for the observed particles. We effectively employed cryoPROS in the cryo-EM single particle analysis of the hemagglutinin trimer, showing the ability to restore the near-atomic resolution structure on non-tilt data. Moreover, the enhanced version named cryoPROS-MP significantly improves the resolution of the membrane protein Na_X_ using the no-tilted data that contains the effects of micelles. Compared to the classical approaches, cryoPROS does not need special experimental or image acquisition techniques, providing a purely computational yet effective solution for the preferred orientation problem. Finally, we conduct extensive experiments that establish the low risk of model bias and the high robustness of cryoPROS.

## 1 Introduction

Recent advancements in cryo-EM hardware and image processing software have ushered in a transformative era in structure determination, establishing it as the prevailing method in the structural biology domain. Notably, Jacques Dubochet’s immersion freezing workflow has become a foundational aspect of cryo-EM technology, demonstrating broad applicability across a range of biological specimens [1, 2]. Yet, challenges remain: factors such as ice quality on the grid and particle distribution within the ice can significantly impact the results. Consequently, preparing high-quality samples continues to be a critical requirement for successful 3D structure determination through single-particle cryo-EM [3].

A main challenge during the stage of frozen samples that impedes structural analysis is the issue of preferred orientation [4–10]. In an ideal scenario, biomacromolecules of interest should exhibit uniformly random orientations within the vitreous amorphous ice. However, it is commonly observed that samples tend to adopt a specific, dominant orientation due to interactions at the air-water or the support-water interface [11, 12]. This phenomenon can significantly degrade the resolution and quality of the resulting map, often leading to misleading density maps known as preferred orientation artifacts. Addressing this preferred orientation issue is widely recognized as an important step in refining cryo-EM as a more universal method for structural analysis [6, 10, 13–17], making it the go-to technique for high-throughput biomolecular research and drug screening [18–20].

Over the years, numerous attempts have been made to confront the issue of preferred specimen orientation in cryo-EM, with most efforts centered around grid preparation and data collection. Techniques such as the use of detergents [15, 21–23], ice thickening [24], and specific biomolecule modifications [25] have been explored. While these methods showed promising results on specific proteins, they were often encumbered by extensive condition screening, added complexities, and time-intensive protocols without consistent success. An alternative proposition suggested using hydrophilic or selectively capturing modified graphene as a support to standardize the particle pose distribution [26, 27]. However, the preparation and modification of graphene proved labor-intensive and achieved limited success in fully resolving the orientation problem. In addition, compensatory techniques during data collection, particularly tilting strategies [7], were introduced. These methods bypassed the challenges of sample preparation but introduced other drawbacks, such as decreased image acquisition efficiency, escalated beam-induced movement during tilting, the imperative of precise defocus gradient estimation, and increased ice layer thickness due to geometric constraints [12]. For small molecules, aggregation complications also arose, somewhat limiting the effectiveness of the tilting methodology. Despite these difficulties, tilting is currently one of the most effective solutions to tackle preferred orientation in cryo-EM studies. Given the aforementioned limitations in addressing the preferred orientation during the wet-lab phase, designing the computational approach becomes an attractive alternative, which motivates our work.

We first conduct extensive experiments demonstrating that the main challenge in addressing the preferred orientation issue lies in the misalignment of particle images during the refinement process due to unevenly distributed signals from different views. This inaccurate orientation estimation leads to poor reconstruction quality, particularly impacting the axial resolution. In contrast, the missing wedge issue in computational tomography (CT) lacks projected data from certain view angles; however, the corresponding orientation information for the observed particles is always accurately known. Therefore, equating the missing wedge issue in CT with the reconstruction artifacts observed in cryo-EM single particles exhibiting preferred orientation is a misconception. Such conflation obscures our accurate understanding of the problem. Sorzano *et al.* have posited this claim [28–30], and we further validate its soundness.

Motivated by the above observation, we introduce cryoPROS (**PR**eferred **O**rientation **S**olver), an AI-based computational approach designed to address preferred orientation challenges in cryo-EM. Using a conditional variational auto-encoder (CVAE) model, cryoPROS generates high-quality auxiliary particles with diverse orientations, significantly improving the alignment accuracy of the observed particles. As a pure computational framework, cryoPROS does not require specialized sample preparations, specific data acquisition methods, or further computational refinement, substantially reducing experimental complexities. Our experimental results demonstrate that cryoPROS can obtain the near-atomic resolution structure using the highly preferred oriented dataset of hemagglutinin (HA) trimer (EMPIAR-10096). Furthermore, for the samples that are embedded in detergent micelles or lipid nanodiscs, we propose a variant of cryoPROS named cryoPROS-MP. This enhanced version has demonstrated a significant resolution improvement over conventional methods using the non-tilted data of the membrane protein Na_X_. Finally, we test various configurations that confirm the low model bias and the high consistency of cryoPROS. This makes cryoPROS an effective and reliable tool, ideally supplementing traditional structural determination methodologies and extending its range of applications.

## 2 Results

We next present a comprehensive analysis of the preferred orientation problem, followed by introducing our novel algorithm, cryoPROS, designed to address this challenge. We illustrate the practical implementation of cryoPROS on multiple datasets and examine its characteristics and performance in-depth.

### 2.1 Orientation estimation bias: the main issue for structure determination from data with preferential orientation

We conducted a series of simulated tests to identify the primary computational challenge associated with the preferred orientation problem. To do this, we synthesized two datasets, called SIM1 and SIM2, by projecting the density map of the HA trimer. The density map was derived from the atomic model (PDB ID: 3WHE) and the projection process was implemented using the relion project module within Relion. We added Gaussian noise to the particles, with a zero mean and a standard deviation of 60. Both SIM1 and SIM2 contain 130,000 particles, and the average signal-to-noise ratio (SNR) is -15.63db. In SIM1, the orientation of particles was sampled from a uniform distribution in SO(3). In contrast, the particles in SIM2 exhibit a preferred orientation, with 82,909 particles oriented near the Z-axis, as shown in (Fig. 1a). We reported the results under the following three procedures:

1. SIM1-Reconstruction: reconstruction of SIM1 using true orientations;
2. SIM2-Reconstruction: reconstruction of SIM2 using true orientations;
3. SIM2-Refinement: reconstruction of SIM2 using orientations estimated by the autorefine module in CryoSPRC.

**Fig. 1.**
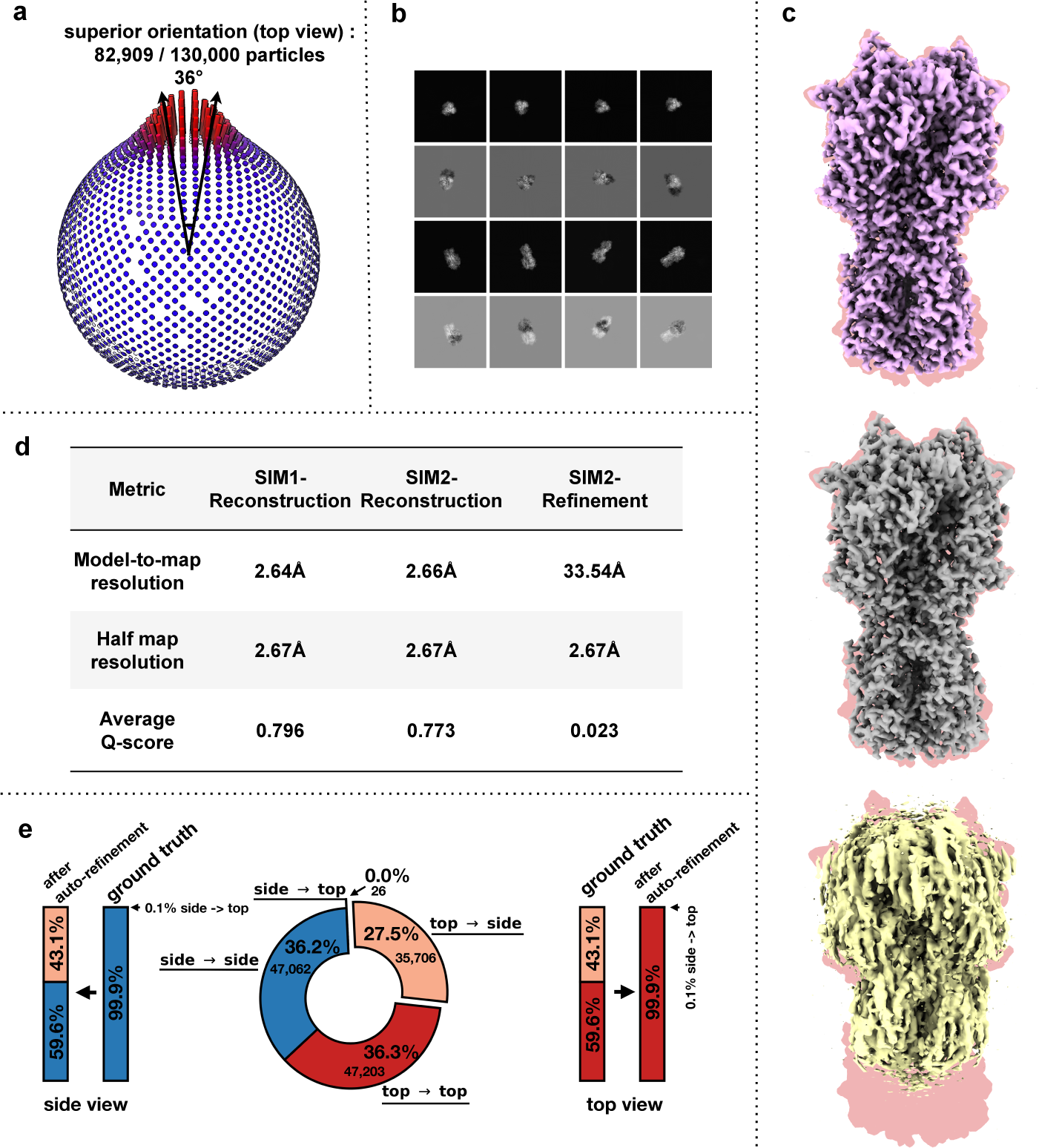
Effect of bias in orientation estimation on determining structures from preferentially oriented data. **a**, Pose distribution of SIM2 dataset, emphasizing the superior orientation defined as being within 36*^◦^* of the Z-axis. The proportion of this superior orientation is indicated. **b**, From the SIM2-Refinement dataset, eight particles were chosen, represented by eight pairs of top-bottom images, resulting in a total of 16 images arranged in a 4 *×* 4 grid. For each pair, the top image shows the ground-truth projection (without added noise) of the HA trimer, while the bottom image represents the difference between the projection (with its orientation estimated by CryoSPARC’s autorefine) and the ground-truth projection. Among these eight particles, the ground truth orientations for half are from the superior orientation (top view), while the other half is from the inferior orientation (side view). **c**, Density maps are presented from top to bottom in the following order: SIM1-Reconstruction (in violet), SIM2-Reconstruction (in gray), and SIM2-Refinement (in yellow). **d**, Quantitative comparison of indicators for the density maps displayed in **c**. **e**, In SIM2-Refinement, CryoSPARC’s autorefine incorrectly estimated 59.6% of the particles from a superior (top view) to an inferior orientation (side view), as shown on the right. These misassignments account for 35, 706 particles or 27.5% of the total, overwhelming the actual side view, as highlighted in the middle. In SIM-Refinement, misassigned top views made up 43.1% of the side view orientation, as illustrated on the left.

The reconstructed density maps are presented in (Fig. 1c). The density maps from SIM1-Reconstruction and SIM2-Reconstruction are similar, exhibiting comparable quantitative performance in terms of FSC-based metrics and Q-score (Fig. 1d). However, the density map produced by SIM2-Refinement shows significant deterioration, particularly in the loss of axial density due to the preferred orientation (Fig. 1c and Fig. 1d). This degradation appears to be induced by the misalignment (Fig. 1e) during the refinement process. In SIM2-Refinement, approximately 43.1%, 35, 706 out of 82, 709, of the particles with a superior orientation (top view) were incorrectly estimated as having an inferior orientation (side view) by CryoSPARC’s autorefine (Fig. 1e, right). As a result, these incorrectly assigned particles of the top view, constituting around 43.1% of the side view orientation (Fig. 1e, left) in the reconstruction stage, significantly impacted the resolution.

The above observations motivated us to re-evaluate the challenges posed by the data with preferred orientations. The conventional approach to single-particle cryo-EM structure determination is an iterative process, starting with particle alignment through projection matching and followed by reconstruction. These steps generate estimated orientations and density maps, respectively. However, this method encounters considerable difficulties when addressing preferentially-oriented datasets. In the reconstruction phase, uneven coverage of the Fourier space results in reconstructions with anisotropic resolution. Furthermore, when a density map from the previous iteration is employed as a refinement reference, it inevitably leads to misalignment. This misalignment introduces orientation estimation errors, further reducing the quality of the reconstruction. As this iterative process continues, orientation estimation errors accumulate, leading to an increasing bias. This accumulating bias either progressively deteriorates the density or results in entirely erroneous densities. When overfitting occurs, it accentuates this decline, further undermining the reconstruction quality. In summary, the challenges of preferred orientation in single-particle cryo-EM primarily stem from orientation estimation bias. Additionally, we expanded our investigation to include real data, as detailed in Section 2.5. These findings further supported the above analysis related to datasets with preferred orientations and inspired us to explore computational approaches that can accurately determine the poses of particles.

### 2.2 The cryoPROS method

We introduce cryoPROS, an AI-based method designed for accurate pose estimation in preferentially oriented data. Specifically, cryoPROS consists of two main modules: the generative module and the refinement module, as illustrated in Fig. 2. The generative module leverages a Conditional Variational Autoencoder (CVAE) framework [31, 32], taking as its inputs a highly preferentially oriented raw particle stack, estimated imaging parameters, and a 3D latent volume. By minimizing the conditional evidence lower bound, we establish a self-supervised loss that eliminates the need for an additional supervised dataset (see Methods). After training, the network can generate auxiliary particles with uniformly distributed orientations. The refinement module combines the raw particles with the generated particles and employs cryo-EM single-particle analysis software (e.g., Relion, cryoSPARC, or cisTEM) to perform ab initio reconstruction and pose estimations. It is worth mentioning that the generated particles are only used for estimating poses, not for reconstructing the density map.

**Fig. 2.**
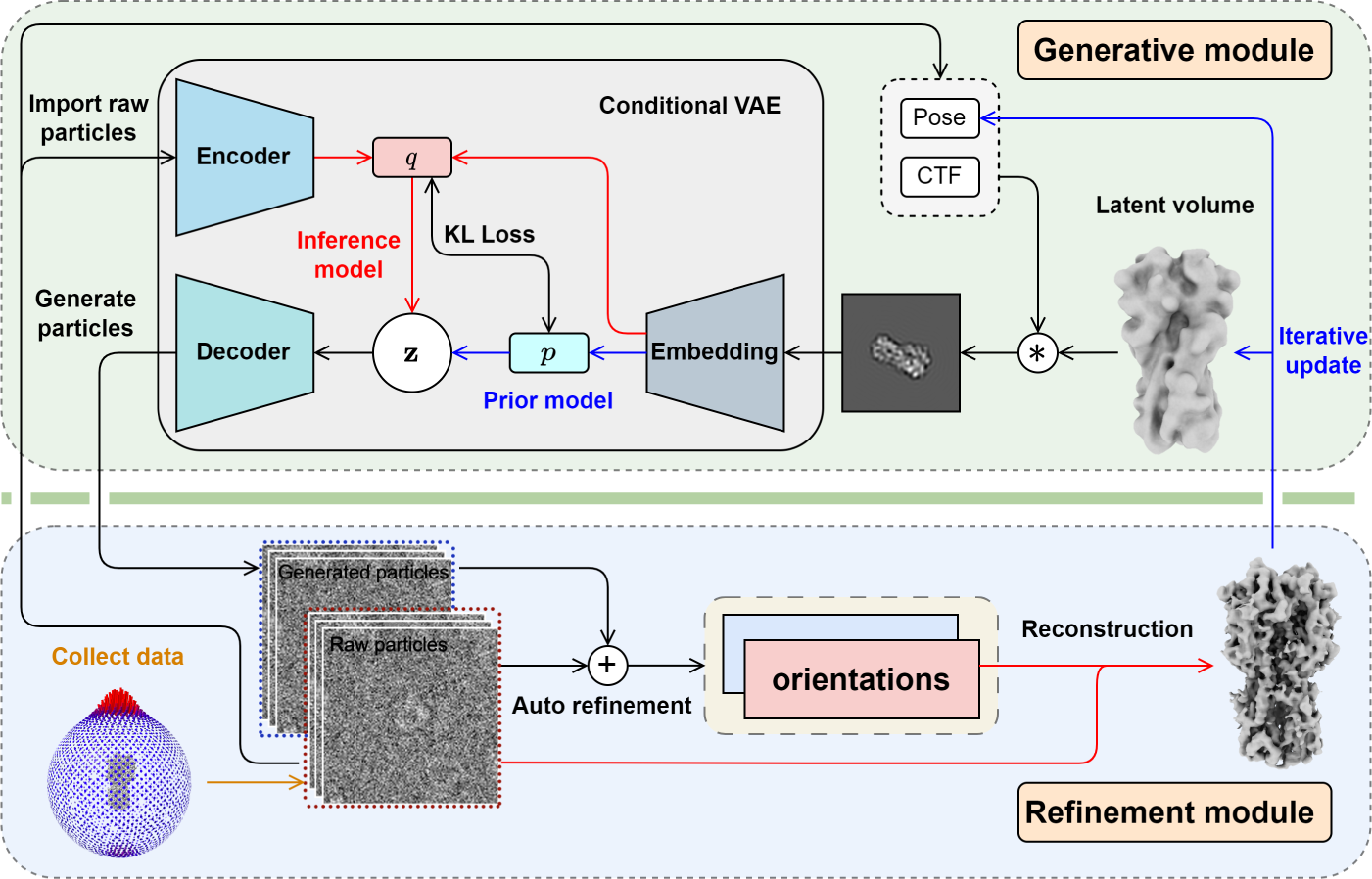
The cryoPROS method for single-particle cryo-EM preferred orientation problem. The overall protocol of cryoPROS contains generative and refinement modules, applied iteratively. The generative module employs a conditional variational autoencoder (CVAE) framework, incorporating a latent volume in the latent space and integrating a physical imaging process. It generates projection particles with predetermined imaging parameters after training. The refinement module combines raw particles with generated particles, utilizing cryo-EM single-particle analysis software for *ab initio*orientation estimation. The reconstruction employed only the raw particles, omitting the generated ones, and used the updated orientations to produce a more precise 3D density map.

The CVAE model in cryoPROS consists of two components: the conditional encoding stage and the decoding stage. In the conditional encoding stage, we integrate the encoding features of both the raw particle and the latent particle using given imaging parameters (CTFs, poses) and a latent volume to form an inference model *q* for the latent variable **z**. Sampling from the inference model *q*, the decoding network maps the latent variable to a synthesized particle, thereby generating a reconstruction loss in relation to the input raw particle.

Moreover, we directly encode the latent particle to obtain a prior model *p* and impose a loss based on the Kullback-Leibler (KL) divergence between the inference model *q* and the prior model *p*. After training, we can generate auxiliary particles by decoding samples from the prior model *p*, inputting the inferior orientations and CTF parameters. To enhance its expressive ability, we designed a hierarchical VAE structure [33, 34] (see Methods and Supplementary Section 6).

Challenges arise when target proteins are situated in micellar environments not present in the homologous protein or associated with different micellar contexts. In such cases, reconstructing the relevant micellar environment for the low-pass filtered homologous protein, used as the initial latent volume, becomes crucial. Therefore, we propose cryoPROS-MP, which is customized for membrane proteins in detergent micelles or lipid nanodiscs. Compared to cryoPROS, cryoPROS-MP includes an additional procedure for reconstructing the micellar information (see Methods).

### 2.3 Validation with several preferential oriented datasets

We conducted validation of cryoPROS on SIM2 and three experimental datasets: the PO-subset of TRPA1, obtained by removing side view particles from the TRPA1 dataset (EMPIAR-10024); the non-tilt HA trimer dataset (EMPIAR-10024); and our uniquely collected Na_X_ dataset. The relevant details of the data and the corresponding parameters can be seen in Section 1 and Table 1 in the supplementary material. In these experiments, the preprocessing, including motion correction and particle-picking, adhered to the standard cryo-EM SPA workflow. Additionally, the refinement module utilized the widely-used cryo-EM orientation estimation software, CryoSPARC, with its default settings.

Two iterations of cryoPROS were carried out. In the first iteration, the homologous protein’s density map was lowpass filtered to 10^Å^, serving as the starting point for the latent volume. The latent volume is updated using the reconstructed density map from raw particles using estimated orientations from the first iteration. It is shown in Section 2 and Fig. 1 in supplementary material that the impressive fidelity of cryoPROS-generated particles in simulating the signal of raw cryo-electron microscopy (cryo-EM) data. Additionally, we employed post-processing methods, such as EMready [35], as an optional step to address artifacts arising from uneven orientation. We measure the quality of the reconstructed density using the half-map resolution, model-to-map resolution, and Q-scores. These measures collectively serve to validate the density maps generated through cryoPROS quantitatively.

### 2.4 SIM2: eliminating orientation estimation bias

We applied cryoPROS to the SIM2 dataset by choosing a homologous protein (PDB ID: 2RFU) with 16% sequence identity to the target protein as the latent volume. The results demonstrated substantial improvements in the density map, particularly in effectively restoring the lost density along the Z-axis. A model-to-map resolution of 3.28^Å^ was achieved by cryoPROS (Fig. 3a, in violet), as determined by the HA trimer atomic model (PDB ID: 3WHE). This represents a significant improvement compared to the resolution of 33.54^Å^ obtained from conventional auto-refinement (Fig. 3a, in yellow).

**Fig. 3.**
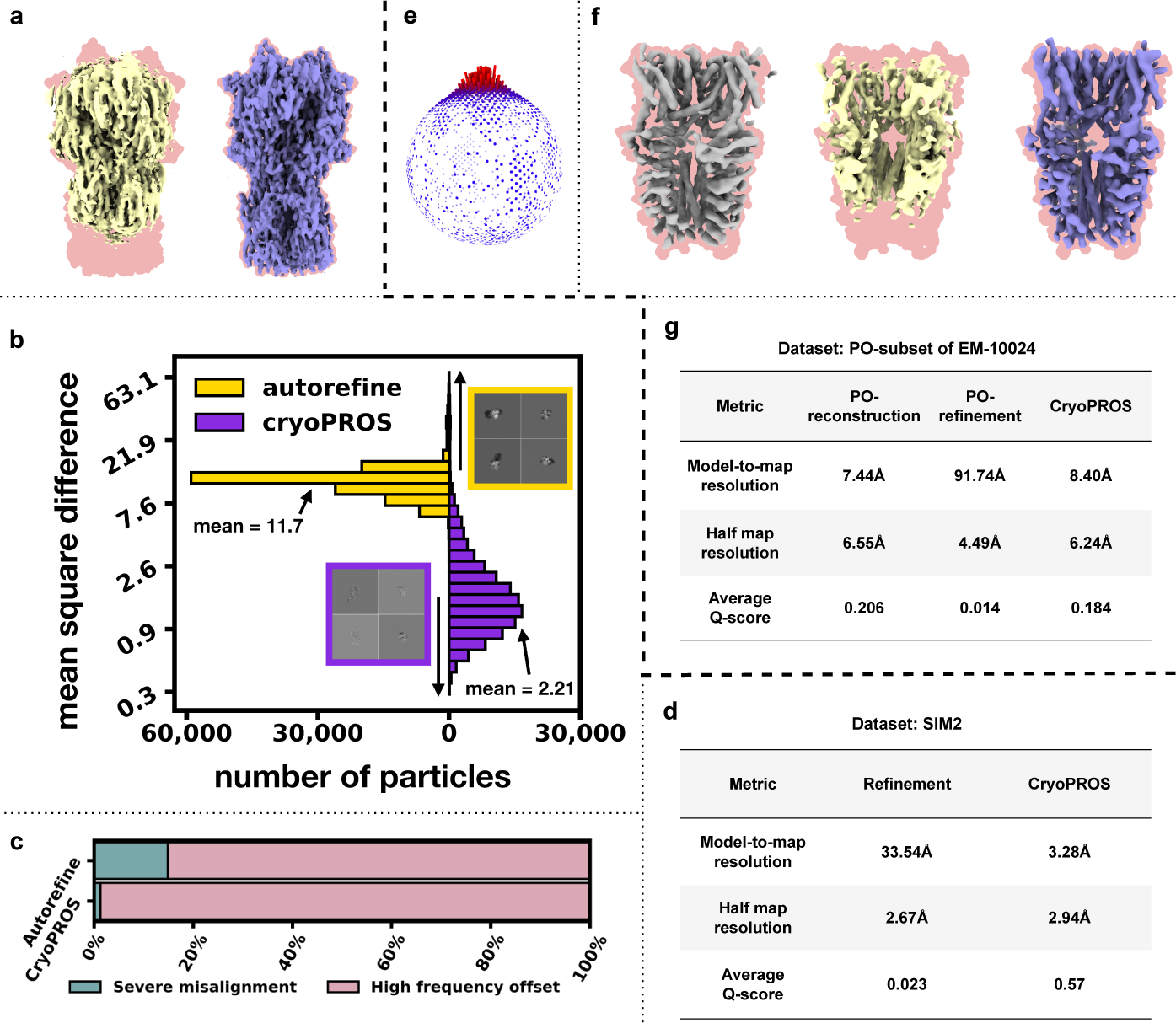
Avoiding orientation estimation bias with cryoPROS. **a**, Reconstructed density maps of SIM2 dataset via autorefinement (in yellow) and cryoPROS (in violet). **b**, Histograms of MSE the noise-free ground truth projections and the projections derived from the orientation estimated by cryoPROS and CryoSPARC’s autorefine, respectively. Projection differences were highlighted by boxes of yellow and violet, respectively. Means of the MSEs were labeled. **c**, Proportion of severe misalignment (in green) and high-frequency (in red) offset in autorefinement and cryoPROS misalignment. **d**, FSC-based resolutions and Q-scores of maps in **a** are listed. **e**, Pose distribution of constructed PO-subset of TRPA1. **f**, Reconstructed density maps of PO-subset of TRPA1 via reconstruction only from fixed pre-estimated and more accurate orientations (in gray), autorefinement (in yellow), cryoPROS (in violet), respectively. Density maps in **a** and **f** were superimposed on the envelope of their ground truth atomic models (in red). **g**, FSC-based resolutions and Q-scores of maps in **f** are listed.

We further compared the pose assignment accuracy between cryoPROS and conventional auto-refinement by computing the mean square difference (MSE) between the noise-free ground truth projection and the projection derived from the orientation estimated by CryoSPARC’s auto-refine (Fig. 3b). Reduced MSEs of residuals from cryoPROS were observed, indicating overall higher orientation accuracy. Using residuals greater than 13 as the threshold for significant orientation errors, cryoPROS decreased the rate of severe misalignments from 14.85% to 1.27% (Fig. 3c). This led to enhanced resolution (Fig. 3d). These results show the proficiency of cryoPROS in addressing misalignment challenges and achieving superior resolutions.

### 2.5 PO-subset of TRPA1: restoring missing density

We assessed cryoPROS’s ability to restore missing density caused by orientation estimation bias in the case of TRPA1’s constructed preferred-oriented dataset. TRPA1 is a 690kDa prototypical membrane protein and acts as a detector for toxic chemical agents encountered in the environment or produced during tissue damage or drug metabolism [36].

We constructed a subset from the deposited raw particle images (EMPIAR-10024 [36], containing 43,585 particles) by selecting all top views and a limited number of side views. This resulted in a preferentially oriented subset, referred to as the PO-subset, containing 14,436 particles. The pose distribution of this subset can be found in Fig. 3e. Assuming accurate alignment of the raw particle images, we retained the estimated orientation and reconstructed the PO-subset. This process generated a density map, which we named the PO-reconstruction. Despite a slight drop in resolution due to a reduced number of particles, the map remained devoid of artifacts related to preferred orientation (Fig. 3f, in gray), presenting a holistic structure.

In constrast, the traditional refinement applied to the PO-subset, referred to as the PO-refinement (Fig. 3f, shown in yellow), resulted in an unsatisfactory density map. This map suffered from significant density loss along the preferred orientation axis. Furthermore, its quantitative indicators, such as Q-scores and FSC-based resolutions, were notably lower compared to those of the PO-reconstruction (Fig. 3g). These observations demonstrate the effects of orientation estimation bias when analyzing structures in preferentially oriented data.

Next, we applied cryoPROS to the PO-subset. Since no deposited structures of homologous proteins were available, we used Alphafold to predict the rat-derived TRPA1 protein (UniProtID: F1LRH9). After the prediction result was lowpass filtered, it served as the first-round input for cryoPROS. The sequence identity between the predicted rat TRPA1 and TRPA1 was 77%. After two iterations, cryoPROS generated a more accurate and complete density map of TRPA1 (see Fig. 3f, right, violet), achieving resolutions of 6.4Å and 8.4Å based on the half-map and map-to-model FSC criteria, respectively (Fig. 3g). Furthermore, the Q-scores of the density map showed significant improvement (Fig. 3g). CryoPROS’s output resembled the results obtained from PO-reconstruction, indicating its resistance to the density loss caused by the preferred orientation.

### 2.6 HA Trimer: achieving near-atomic resolution structure using non-tilted data

We utilized cryoPROS to process the non-tilt data with preferential orientation sourced from the EMPIAR repository (EM-10096) and conducted an extensive comparison with the results derived from data using the tilt technique. The tilt data was also obtained from the EMPIAR repository (EM-10097). For the initial round of cryoPROS, we selected a homologous protein (PDB ID: 6IDD) with 47% sequence identity to the target protein for lowpass filtering as the latent volume. CryoPROS produced uniformly oriented particles after two iterations, with the pose distribution displayed at the bottom of Fig. 4c. The refinement module then yielded complete and high-resolution results (Fig. 4a, illustrated in blue). Visually, the quality of the density showed substantial improvement compared to the automatic refinement on non-tilt data, and outperformed the automatic refinement on 40*^◦^* tilted data (Fig. 4b, depicted in pink). However, it is slightly inferior to the state-of-the-art results on tilted data (Fig. 4b). Achieving such results necessitates complex subsequent refinements at the per-particle level, including multi-round 3D classification, defocus refinement, and Bayesian polishing [37]. In addition, we post-processed the cryoPROS result using EMReady to eliminate the preferred orientation artifact. This post-processing led to a minor improvement in the quality of results (Fig. 4a, depicted in blue). Throughout the entire process, we refrained from filtering particles based on experience, thereby ensuring a faithful representation of the non-tilt data.

**Fig. 4.**
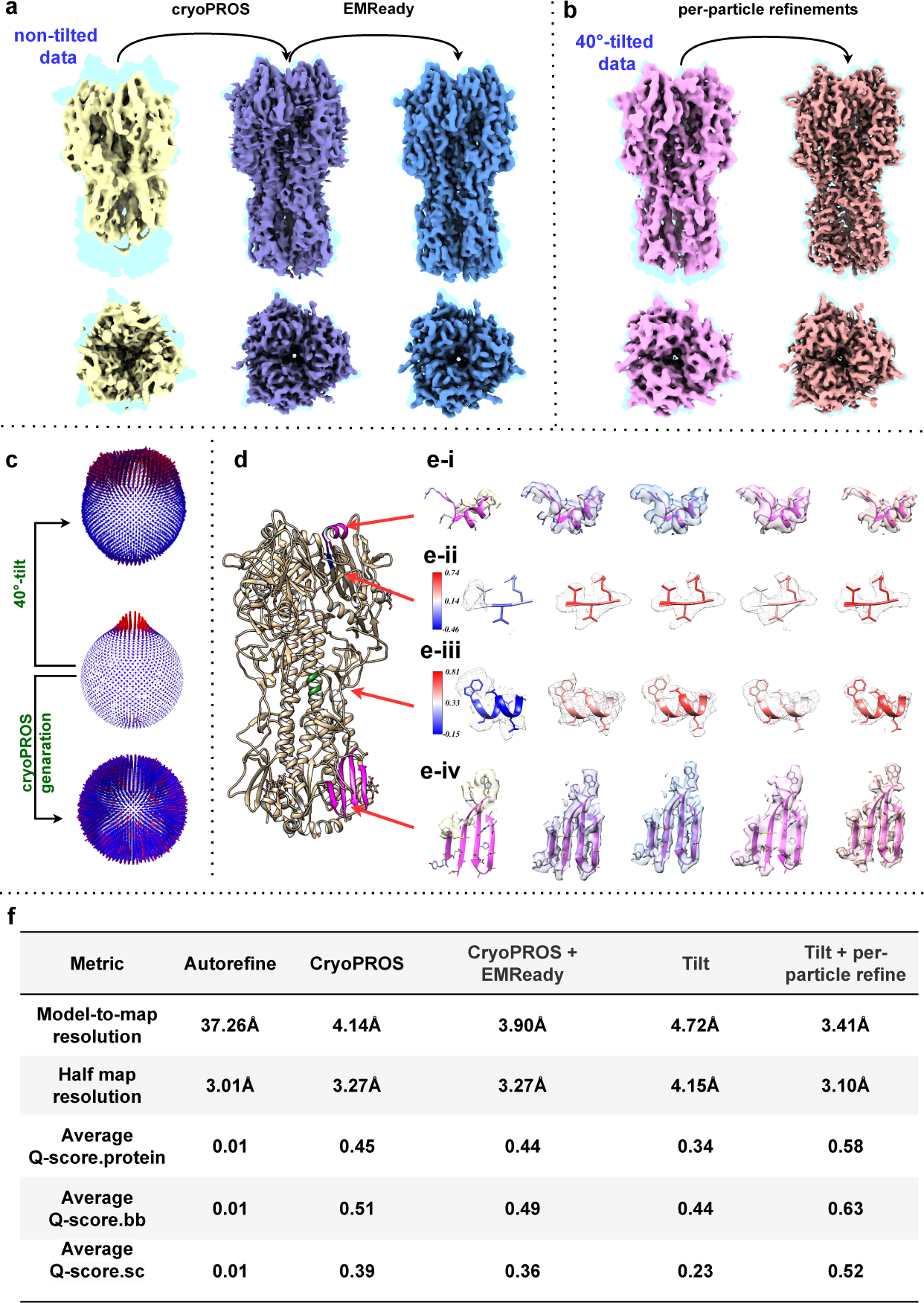
CryoPROS enables the recovery of near-atomic-resolution information from highly preferentially oriented data of the influenza A HA-trimer. **a**, Reconstructed density maps of the non-tilt dataset using: autorefinement (in yellow), cryoPROS (in violet), and cryoPROS with follow-up EMReady postprocessing (in blue). **b**, Reconstructed density maps of the tilt-collected dataset: Autorefinement (in pink) and state-of-art result (in crimson). Notably, achieving the state-of-the-art result necessitates intricate subsequent refinements at the per-particle level, which encompass multi-round 3D classification, defocus refinement, and Bayesian polishing. Maps in **a** and **b** are superimposed on the envelope of the HA-trimer atomic model (PDB ID: 3WHE, in light blue). **c**, Pose distribution of datasets: 40*^◦^*-tilted dataset (top), non-tilt dataset (middle), and cryoPROS generated auxiliary particles (bottom). **d**, Colored parts are selected for close-up in **e**. **e**, Detailed close-ups of selected parts of the density maps shown in **a** and **b**, of which order are consistent with. Panel e-i,iv show the regions of alpha-helix and beta-sheet in low transparency. Panels ii and iv show the selected regions in gray mesh style, embedded atomic model is colored by average Q-score.sc value and average Q-score.bb value, respectively. **f**, Multiple indicators are used to quantitatively comparel density maps shown in **a** and **b**. Q-score evaluations span the atomic level, main chain level, and side chain level, termed as average Q-score.protein, average Q-score.bb, and average Q-score.sc, respectively.

We assessed the quality of density maps using two quantitative metrics: resolution and average Q-score. The resolution measures included the half-map and map-to-model resolutions relative to the HA trimer structure (PDB ID: 3WHE). Q-score evaluations were conducted at the atomic level, main chain level, and side chain level (Fig. 4f). Notably, the cryoPROS output, following post-processing, demonstrated an impressive model-to-map resolution of 3.90 Å and a half-map resolution of 3.27 Å. This result validates the effectiveness of cryoPROS in refining preferred orientation data to nearly atomic resolution. Significantly, this achievement was realized without necessitating additional data collection or the complex per-particle operations typically used to mitigate the intrinsic limitations of the tilt strategy.

Furthermore, we selected specific sites in the known atomic model (PDB ID: 3WHE) for density comparison (Fig. 4d,e). The results demonstrated that the density map generated by cryoPROS was well-preserved, displaying distinct regions with alpha-helical pitch, beta-strand separation, and bulky side chains (Fig. 4e-i, e-ii, shown in blue). We employed Q-score.bb (backbone, see Fig. 4e-iii) and Q-score.sc (side chain, see Fig. 4e-iv) to color the atomic model, which revealed high scores for both the main chain and side chain in the results produced by cryoPROS. In all comparisons, cryoPROS closely mirrored the results derived from the analysis of tilted data. Moreover, in most instances, cryoPROS surpassed the results from tilt-collection-autorefinement (Fig. 4b, depicted in pink), nearly matching the best results from the current dataset. All these findings suggest that cryoPROS can achieve nearly atomic-resolution density maps from preferentially oriented single-particle datasets.

### 2.7 Na_X_: Expanding cryoPROS’s applicability to membrane protein analysis

Na_X_ is an atypical sodium channel involved in regulating sodium homeostasis and other physiological functions, distinguished by its voltage-insensitivity and resistance to tetrodotoxin. It is a potential target for diseases related to sodium balance and other associated conditions [38]. To study this, we gathered a non-tilt dataset containing 411, 823 particle images. Since Na_X_ is a membrane protein, it was purified and solubilized using the LMNG detergent for this dataset. This process results in the detergent’s visualization as a micelle. Particles in this dataset display a strong preferred orientation, rendering side views or inferior orientations nearly invisible. Furthermore, since they are solubilized by detergent, the sizable micelle presence further complicates orientation estimation. Such disordered densities cannot be overlooked. Thus, we applied cryoPROS-MP, an enhanced version of cryoPROS, to this case.

We used the density map derived from the atomic model of Na_v_1.6 (PDB ID: 8FHD) and applied a lowpass filter to 10^Å^. This map served as the latent volume in the initial iteration of cryoPROS-MP. It shares a 56% sequence identity with the ground truth.

Distinct from cryoPROS approach, cryoPROS-MP’s primary enhancement includes a micelle reconstruction step. This captures the latent volume embedded within the micelle, as depicted in pink in Fig. 5d. We explore the significance of micelle reconstruction to cryoPROS-MP in greater depth in Supplementary Section 3 and Supplementary Fig.2. When juxtaposed with the conventional autorefinement result (shown in yellow in Fig. 5b), the outcome from cryoPROS-MP (displayed in blue in Fig. 5b) demonstrates a marked improvement in map quality, particularly emphasizing the clarity of the central transmembrane regions. The atomic model, colored based on the backbone-level average Q-score of the cryoPROS-MP result, highlights the congruence between the map and the model, especially in the central areas (refer to Fig. 5d). However, some relatively flexible exterior regions show a less consistent alignment. Using the ground truth atomic model as a benchmark, the cryoPROS result attains a model-to-map resolution of 7.22Å, with additional validation indicators presented in Fig. 5c. Though this resolution is within the medium spectrum, it represents a significant leap over conventional refinement techniques. This resolution constraint largely stems from the total lack of data orientation. In conclusion, cryoPROS-MP serves as an effective expansion of the cryoPROS framework, specifically tailored to membrane proteins. Considering their crucial role in drug discovery, we emphasize the indispensable nature of this enhancement.

**Fig. 5.**
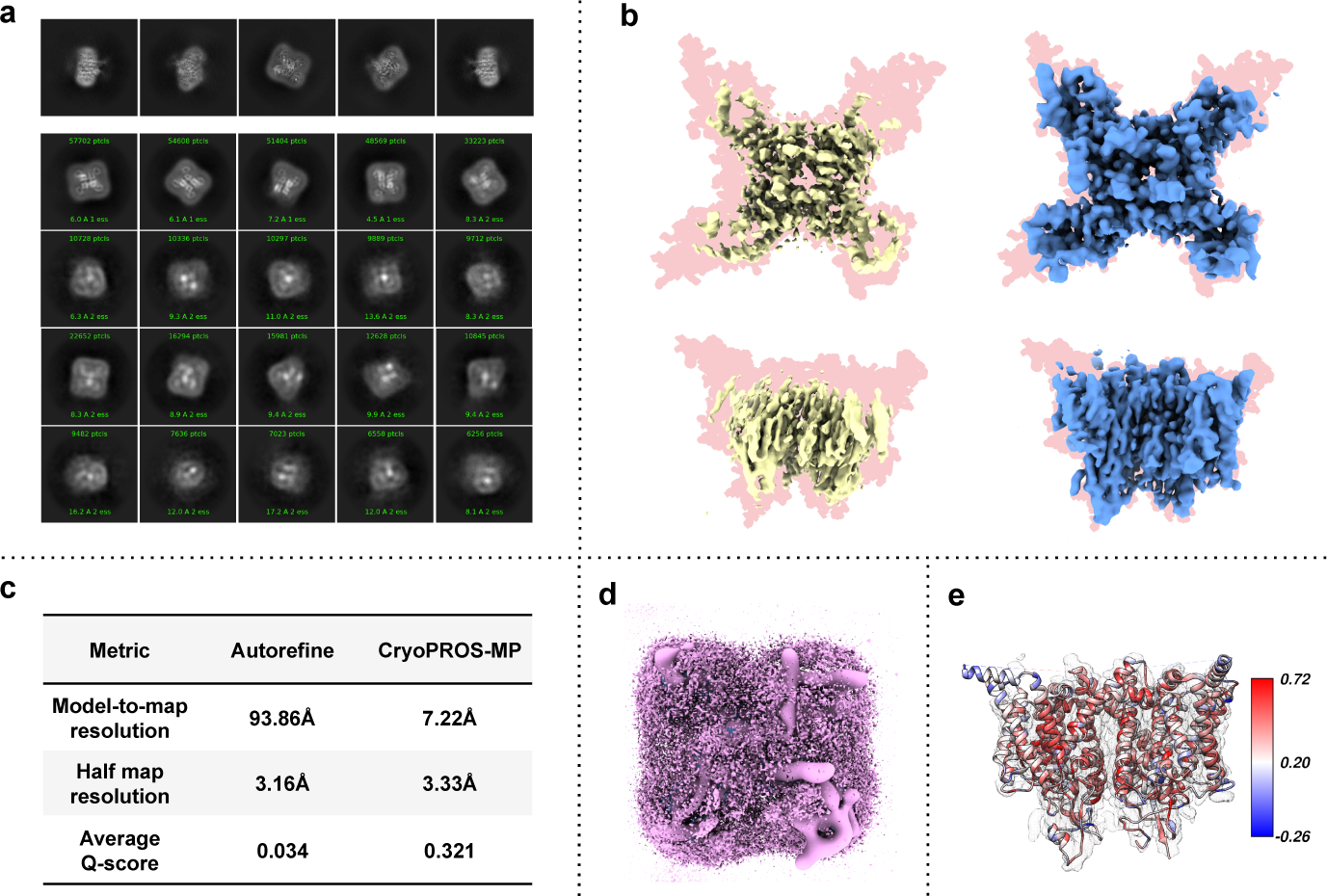
CryoPROS-MP: Adapting to membrane proteins solubilized in micelles or liquid nanodiscs. **a**, The upper row displays projections of the ground truth density map, with orientations chosen at random. The 2D classification averages of the Na_X_ dataset, obtained via CryoSPARC, are arranged in a 4 *×* 5 grid below. **b**, Density maps reconstructed from the non-tilt data using autorefinement (in yellow) and cryoPROS-MP (in blue) are presented. These maps are superimposed on the envelope of the ground truth atomic model (in light red). The side view is displayed in the upper row, while the top view occupies the lower row. **c**, FSC-based resolutions and Q-scores of maps in **b** are listed. **d**, Latent volume for the first iteration of cryoPROS-MP. **e**, The ground truth atomic model is colored by the backbone-level average Q-score of the resulting cryoPROS-MP map. This map, after being made partially transparent, is superimposed onto the atomic model.

### 2.8 CryoPROS is unaffected by the potential issue of model bias

Given that cryoPROS requires a lowpass filtered map of a homologous protein as an initial volume for the first iteration, we examined the potential for model bias. Since cryoPROS requires a low-pass filtered map of a homologous protein as an initial volume for its first iteration, it is crucial to evaluate whether model bias influences the resulting map from CryoPROS. CryoRPOS implements three strategies to mitigate this risk.

In its first strategy, cryoPROS undergoes two rounds of iterations. After the first iteration, cryoPROS updates the latent volume, which is reconstructed exclusively from raw particles while excluding the generative ones. Consequently, no orientation estimation directly relies on the projections derived from the homologous protein. Therefore, the risk of model bias is reduced.

In its second strategy, during the training process, the CVAE employs a reconstruction loss that aligns the generated auxiliary particles with raw particles. This step eliminates any detailed features of the homologous protein that might misguide the reconstruction of the target protein. We discovered that the noise generated by the trained CVAE is highly unlikely to overfit into the density maps. In comparison, Gaussian noise is susceptible to overfitting, a phenomenon widely recognized as the “Einstein from noise” effect [39]. To further explore this, we conducted the “Einstein from noise” experiment. This involved parallel autorefinement of both Gaussian pure noise stacks and CVAE-learned pure noise stacks. We utilized varying degrees of lowpass filtered density maps as references (specifically the ribosome, with PDB ID: 6pcq). The resulting three-dimensional signal density map is essentially a misleading signal derived from noise due to inherent model assumptions. The finer the resolution, the higher the potential for model bias. As depicted in Fig. 6b, with a density map lowpass filtered to 7^Å^ serving as a reference, CVAE effectively curbs the interference of model bias. Conversely, Gaussian noise demands more extensive lowpass filtering to achieve a similar bias reduction, as shown in Fig. 6b. Fig. 6c highlights that the density maps generated by CVAE-learned noise have markedly lower average quality—roughly two orders of magnitude less than maps produced by Gaussian noise. A closer look at the induced density map slices in Fig. 6d shows that the CVAE-learned noise is nearly indistinguishable due to its minimal impact from its inferior quality. These tests unequivocally underscore the importance of the cryoPROS generative module in effectively sidestepping model bias.

**Fig. 6.**
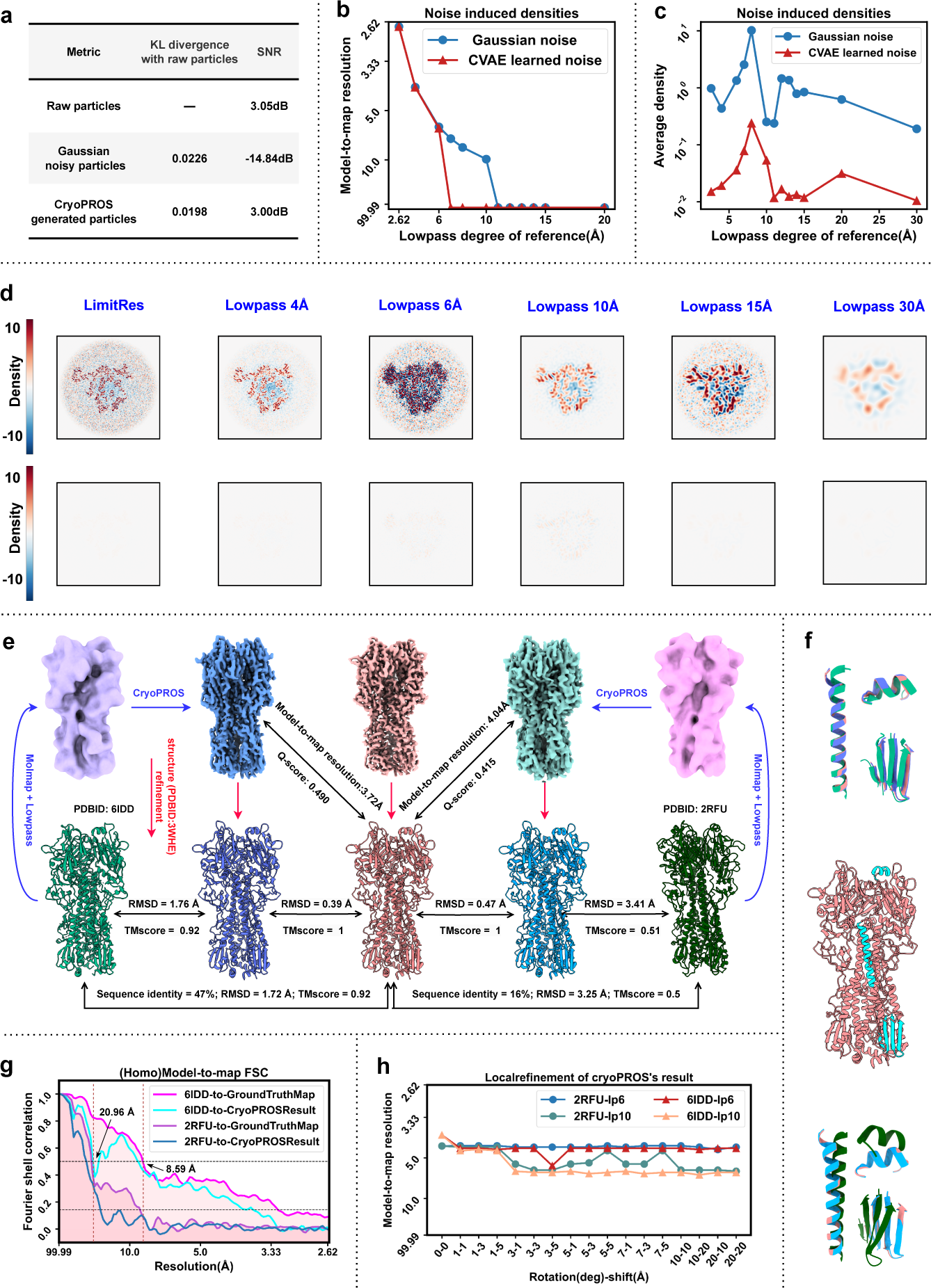
CryoPROS is unaffected by the potential issue of model bias. **a**, Evaluation of the similarity between and generated noisy particles (including CVAE noisy and Gaussian noisy), employing KL divergence and SNR as the evaluation metrics, see Supplementary Section 5.1 for calculation details. **b**, Model-to-map FSC resolution between the alignment reference and the “phantom” density maps obtained from “Einstein from noise” three-dimensional experiment: perform autorefinement on Gaussian pure noise stacks and CVAE pure noise stacks with different degrees of lowpass filtered density maps as references (ribosome, PDB ID: 6pcq). **c**, Comparison of the average density of maps obtained in the “Einstein from nois” three-dimensional experiment. **d**, Slice comparison of the maps obtained the“Einstein from noise” three-dimensional experiment. **e**, CryoPROS experiments using homologous proteins. Two sets of experiments were conducted using homologous proteins different in sequence identities: cryoPROS-6IDD and cryoPROS-2RFU. Density map results (top row) and associated atomic model (bottom row) are presented, along with structure refinements based on the atomic model 3WHE. The homologous proteins utilized in the two experimental groups are introduced on the left and right sides: 6IDD (atomic model shown in light green, lowpass filtered density in light purple); 2RFU (atomic model shown in dark green, lowpass filtered density in pink). The middle three densities are as follows: cryoPROS-6IDD results (displayed in dark blue), best results from the Tilt strategy (shown in red), and cryoPROS-2RFU experimental outcomes (in blue-green), their structure refinement results are shown below, respectively. RMSD and TM-score were utilized to assess similarity and demonstrate cryoPROS’s low model bias risk property. **f**, Alignment and comparison of atomic models. All atomic models in **g** were aligned and compared, focusing on the selected region (middle). The upper and lower sections display relevant results from the two groups of cryoPROS-6IDD and cryoPROS-2RFU experiments, respectively. **g**, (Homologous Protein) model-to-map FSC curve. **h**, Follow-up local refinement experiment on cryoPROS, using different local search parameters, with few changes observed. The two values in the figure’s legend correspondingly signify the homologous protein employed in the cryoPROS experiment and the low-pass parameters utilized in the local refinement experiment. To illustrate, 6IID-lp6 signifies the application of a 6-Å low-pass filter on the 6IID-cryoPROS resulting map as the reference of local refinement and then local search the orientation on the basis of that is 6IID-cryoPROS estimated. The x-axis illustrates the extent of rotation and displacement searches.

CryoPROS generates particles by adding noise to the projections of the latent volume. To simulate this complex process, we adopted the CVAE framework. However, a simpler method to generate particles involves adding Gaussian noise directly to the projections by the latent volume. The detail of this method is given in supplementary Section 5.2. Also, we report the results on HA trimer and Na_X_ in Supplementary Table 3. We found that applying Gaussian noise does not yield satisfactory results for the Na_X_ protein. Although it appears to be effective for the HA trimer, the resultant structure is much closer to the homologous protein than that produced by cryoPROS, suggesting a substantial risk of model bias. This finding validates the fundamental difference between CVAE-learned noise and Gaussian noise. Additionally, we calculated the KL divergence and the noise intensity, as given in Fig. 6a. The results demonstrate that the CVAE-learned noise closely mimics the real noise present in raw particles.

In its third strategy, CryoPROS employs a low-pass filtered initial model derived from a homologous protein that shares only low-frequency information with the target protein, intending to retain only the characteristic signals common to both the homologous protein and the target protein. This methodology is broadly endorsed within the cryo-EM field as a means to circumvent the issue of model bia [30]. For further validation of its effectiveness in the context of CryoPROS, we conducted an ablation study. This study assesses CryoPROS’s robustness against variations in the threshold of the low-pass filter and the sequence similarity between the target protein and the homolog.

We selected a homologous protein with a lower sequence identity (PDB ID: 2RFU, with only 16% sequence identity), denoted as cryoPROS-2RFU. The results of this experiment, illustrated in blue in Fig. 6e, largely align with the previous experiment based on the homologous protein 6IDD (referred to as cryoPROS-6IDD, results shown in blue-green in Fig. 6e). This consistency suggests that the effectiveness of cryoPROS is not heavily reliant on the choice of the homologous protein. The atomic models of the homologous proteins used in these two experiments, as well as the latent volume from the first round of cryoPROS, are displayed in Fig. 6e. In addition, our examination of real data (Na_X_), detailed in Supplementary Section 4.1 and Supplementary Table 2, demonstrates the consistent stability of outcomes when using different homologous proteins. Furthermore, our exploration in Supplementary Section 4.2 and Supplementary Fig.3 delves into the impact of distinct lowpass parameters of the initial latent volume on cryoPROS outcomes. Our findings suggest that cryoPROS maintains a low risk of introducing model bias when various lowpass filter parameters are utilized. However, regarding the accuracy, the current configuration of cryoPROS with a 10Å lowpass is the default choice.

We subsequently compared the results of cryoPROS-6IDD and cryoPROS2RFU with the currently best results (achieved via Tilting+per-particle refinements) for structure refinement based on the atomic model 3WHE. We evaluated the correspondence between the three refined structures using TMscore and RMSD. Both sets of cryoPROS results were found to be remarkably close to the current best results and significantly divergent from the homologous proteins. Similar outcomes were observed in other datasets (refer to Supplementary Section 5.3 and Supplementary Fig.4 for more details). As shown in Fig. 6f, the positional relationship of atomic models from the three groups at different sites confirms that cryoPROS’s results align more closely with the best results procured by tilting, rather than being nearer to the homologous proteins.

Homologous proteins and target proteins are expected to share similar low-frequency features while differing in high-frequency local features. We computed the FSC curves of the atomic model-to-map for homologous proteins, target protein ground truth map, and cryoPROS result, and we determined the resolution based on this FSC curve (see Fig. 6g). Taking homologous protein 6IDD as an example, we found that the correlation between the density map obtained by cryoPROS and the homologous protein is lower than that of the target protein in each frequency band. By setting a threshold correlation of 0.5 (corresponding to 8.59^Å^), we delineated the common characteristic frequency band of the two proteins (lower than 8.59^Å^) and the unique characteristic frequency band of the homologous protein (higher than 8.59^Å^). The correlation of the cryoPROS result fell below 0.5 at 20.96^Å^, which is in the low-frequency band. This suggests that cryoPROS successfully avoids incorporating specific local details from homologous proteins.

Finally, we conducted a series of local refinement experiments subsequent to cryoPROS (see Fig. 6h), which validated the stability of cryoPROS’s results.

## 3 Conclusion and Discussion

We presented cryoPROS, a novel solution that overcomes the key pain point hindering high-resolution analysis of preferentially-oriented data, which is the gradually accumulated orientation estimation bias. CryoPROS corrects this bias through a pure calculation method without the need for additional sample preparation, data acquisition methods, or computational refinement steps, thus reducing the experimental burden. To verify its effectiveness and reliability, we conducted experiments using several datasets and successfully applied cryoPROS to non-tilt data single particle analysis of Na_X_ and HA trimer, achieving high-resolution results comparable to the Tilt strategy. Furthermore, extensive experiments demonstrate cryoPROS’s ability to minimize model bias risk effectively. These findings highlight the significant potential of cryoPROS as a valuable tool for advancing single-particle cryo-EM structural analysis of frozen samples. Moreover, our method encourages open discussions and welcomes further exploration of the following aspects.

### The source of the first round of latent volume

In the standard practice of cryoPROS, the first round of latent volume input is derived from the lowpass filtered homologous protein of the target protein. The homologous protein provides a reliable low-frequency prior to the initial CVAE learning, and its low frequencies are consistent with those of the target protein, effectively improving the quality of CVAE generation. While we have observed cryoPROS’s stable performance with different sequence identities of homologous proteins, the importance of the homologous protein cannot be overlooked. In cases where homologous proteins are lacking, we plan to explore alternative methods, such as predicting an initial model using powerful structure prediction methods like Alphafold2[40], as demonstrated in Section2.5. Another approach is to post-process traditional reconstruction results to obtain a low-pass filtered but relatively complete structure, serving as the first-round input for cryoPROS.

### Potential applications of conditional generative model

In cryoPROS, we have designed a CVAE model capable of generating realistic particles corresponding to any specified pose. The noise distribution in the synthesized particles has been verified to be strikingly similar to that of real noise, presenting a lower model bias risk compared to Gaussian noise. In the field of cryo-EM, both single-particle analysis and electron tomography face the fundamental challenge of managing ultra-low SNR in observed data, thereby emphasizing the critical importance of precise noise modeling. One potential application of this generative model is to utilize the synthesized particles to mitigate the difficulties associated with observed data having imbalanced orientations. For example, in 3D classification, generating particles corresponding to less favored categories could potentially improve the process of particle classification. Moreover, the conditional generative model could be employed to synthesize paired training data for downstream tasks [41]. In signal recovery, a dataset composed of synthesized pairs of clean and noisy particles can be harnessed for training a denoising network, thereby enhancing the signal in the raw particles. Similarly, in pose estimation, the synthesis of particle-pose pairs facilitates the training of a pose estimation network dedicated to determining the orientations of the original particles

### Mixture of preferred orientation and other issues

In real-world scenarios, the preferred orientation problem is often intertwined with other challenges, amplifying the complexity of finding a solution. For instance, membrane proteins embedded in detergent micelles or lipid nanodiscs present additional difficulties, and we have extended the cryoPROS setting to address such cases in Section2.7. More generally, the complexities of cryo-EM analysis lead to a combination of various issues, such as heterogeneity, dynamics, and orientation difficulty in the case of the small protein. Successfully employing cryoPROS to solve these hybrid problems may require further refinement and practical experimentation.

### Post-processing method

In this study, we have divided the preferred orientation problem into two components: orientation estimation bias and anisotropic reconstruction. The core focus of cryoPROS is to address the orientation estimation bias, which is considered more critical. However, in high-resolution structure determination, artifacts resulting from anisotropic reconstruction also will impact structure observation. To address this issue, we introduced EMReady for post-processing the output of cryoPROS, especially in the case of the HA trimer (see Section2.6). Nevertheless, existing post-processing methods like EMReady may not specifically target artifacts related to the dominant orientation density. As a future direction, we aim to develop dedicated tools that can better handle and resolve these artifacts to further enhance the effectiveness of cryoPROS in high-resolution structure analysis.

## Supporting information

Supplementary for Addressing preferred orientation in single-particle cryo-EM through AI-generated auxiliary particles

## Acknowledgements

This work was supported by the National Key R&D Program of China (No.2021YFA1001300) (to C.B.), the National Natural Science Foundation of China (No.12271291) (to C.B.), the Beijing Frontier Research Center for Biological Structure (to M.H.), Shenzhen Medical Academy of Research and Translation (to M.H.) and the National Natural Science Foundation of China (No.12071244) (to Z.S.). We are grateful to Dr. Gaoxingyu Huang for his valuable discussions and to Prof. Yigong Shi for his support in data analysis.

## Data availability

The data that support this study are available from the corresponding authors upon request. Three raw datasets analyzed in this study were downloaded from the EMPIAR repository (EMPIAR-10096, EMPIAR-10096, EMPIAR-10097). SIM1 and SIM2 were generated by the relion project module within Relion. Dataset Na_X_ was collected in-house and will be deposited into EMPIAR in the near future. Structures for the initial latent volume of cryoPROS were downloaded from the Protein Data Bank (PDB ID: 2RFU, 6IDD, 5XL8, 7XM9, 8FHD, 6AGF) or downloaded from AlphaFold Protein Structure Database (UniProt ID: F1LRH9). The structure for validation was based on the HA trimer atomic model (PDB ID: 3WHE) and the TRPA1 ion channel atomic model (PDB ID: 3J9P).

## Code availability

CryoPROS will be open-source upon publication and is also available upon request during the review process.

## Contribution

H.Z, D.Z, C.B., M.H., and Z.S. initiated the project. H.Z., D.Z., C.B. and M.H. developed CryoPROS and carried out testing. H.Z. and D.Z. analyzed the data. Q.W. and N.Y. collected the Na_X_ dataset. H.Z., D.Z., C.B., M.H., and Z.S. wrote the manuscript.

## Competing interests

All other authors declare no competing interests.

## Methods

### The Generative module

The generative module using the CVAE model aims to generate realistic particles, which is the key component in cryoPROS. Let 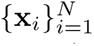 be the set of raw particles, we can obtain the estimated orientation parameters 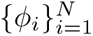 and CTF parameters 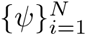 from cryoSPARC and adopt the imaging model as

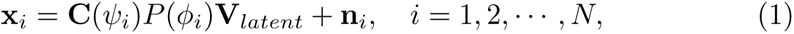

where **C** is the CTF operator depends on *ψ_i_*, *P* (*ϕ*) represents the projection operator for the pose *ϕ*, and **n***_i_* represents the noise. It is noted that the noise term **n***_i_* contains both random error and the model error induced by the estimation error of parameters. Define **Θ** = *{ψ, ϕ,* **V***_latent_}* to be the set of imaging parameters, the CVAE model learns a generative model that maximizes the conditional likelihood *p*(**x***|***Θ**), which is equivalent to the unpaired data modeling problem [41] between the latent projection **v** = **C**(*ψ*)*P* (*ϕ*)**V***_latent_* and raw particles.

Since *p*(**x***|***Θ**) is difficult to optimize directly, the CVAE model involves maximizing a lower bound of log *p*(**x***|***Θ**), known as the conditional Evidence Lower BOund (cELBO). The negative cELBO is given by

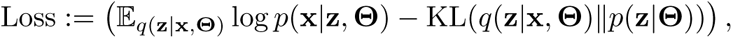

forming the loss function to train our model. Here, **z** is the latent variable, *q*(**z***|***x**, **Θ**) is the inference model, *p*(**z***|***Θ**) is the prior model, and *p*(**x***|***z**, **Θ**) is the generative model. All these models are parametrized by deep neural networks. After the training stage, inputting an imaging parameter **Θ**, the CVAE model samples the latent variable **z** through the prior model *p*(**z***|***Θ**), and subsequently, **z** is transformed into a synthetic particle through the generative model *p*(**x***|***z**, **Θ**). Furthermore, to enhance the representation capability of the traditional VAE model, a hierarchical structure is employed to address the issue of generating unrealistic and blurry samples in the single-layer model. In particular, we assume the latent variable has *L* stochastic layers: **z** = (**z**^1^*,…,* **z***^L^*), and a top-down structure [42] is adopted for the inference and generation process in the CVAE model. Please see Section 6 in the supplementary material for the detailed network architecture.

The CVAE model undergoes training for 20,000 iterations using the Adam optimizer [43]. We set the learning rate to 1 *×* 10*^−^*^4^ and the batch size to 8. The model architecture includes 15 hierarchical layers. To mitigate the issue of posterior collapse, we employ the KL annealing method [44], implementing a linear annealing scheme during the first 10,000 iterations.

### The refinement module

After the generative module, the generated particles are fed into a second module: the refinement module, which aims to optimize the 3D density map reconstruction of preferred orientation data. Generated particles are used for balance pose distribution, helpful for each round of the alignment of the particles, that are of core for structure determination of preferentially oriented single particle samples but problematic to contribute positively in conventional refinement protocol. Specifically, a two-step operation on any popular user-chosen single-particle cryo-EM software, without additional programs: perform autorefinement on a combination of raw and generated particle stacks; then reconstruct raw stack only within fixed refined orientations, obtaining refined pose parameters and 3D density map finally, all of them will be input generative module for improving the generated particles quality and minimize the model bias introduced by homologous proteins to some extent.

It is worth mentioning that all datasets analyzed in this paper consist of preprocessed particle stacks. The initial orientation parameters needed for the first round of cryoPROS can be derived by conducting standard initialization reconstruction and autorefinement on the particle stack, in which intricate preprocessing of raw cryo-EM data is not required. To ensure a thorough understanding, we offer a concise overview of the conventional single-particle preprocessing method. This will enable readers to begin with raw data processing and seamlessly apply cryoPROS in their research. The preprocessing steps consist of several key components. First, apply MotionCor2 to correct the beam-induced motion in the micrograph movie stacks. Next, determine the contrast transfer function parameters for each micrograph by GCTF. Then employ CryoPARC for automatic particle picking in micrographs, and extract the particle stack based on the picked particle coordinates. Subsequently, perform a 2D classification in cryoSPARC, and select well-performing 2D-averaged particles for initial model generation and refinement. Only one optional software/method is introduced here, as others yield similar effects and won’t be detailed further.

## Notes

### Competing Interest Statement

The authors have declared no competing interest.

