## Supplementary for Addressing preferred orientation in single-particle cryo-EM through AI-generated auxiliary particles for "Addressing preferred orientation in single-particle cryo-EM through AI-generated auxiliary particles": Supplementary_230926.pdf

#### 1 Dataset information

In this study, we evaluate the performance and attributes of the newly developed cryoPROS method, utilizing a diverse set of datasets that include both simulated and real-world cryo-electron microscopy (cryo-EM) data. Notably, the SIM2, PO-subset, HA trimer, and $Na_X$ datasets present challenges associated with preferred orientation issues. A detailed evaluation of cryoPROS on these specific datasets is thoroughly discussed in Sections 2.3-7, shedding light on its effectiveness in various scenarios. Comprehensive details of all datasets are carefully cataloged in Table 1, which includes pertinent information such as data source, particle count, target protein, and the resolution achieved by cryoPROS, among other factors.

### 2 Particles generated by cryoPROS

The efficacy of CryoPROS relies on leveraging the powerful representational capabilities of deep learning, specifically through the use of the Conditional Variational Autoencode (CVAE) model for training and generating particle data. This strategic approach aims to rectify the distribution of particle poses within datasets that have a preferred orientation, thereby facilitating the execution of the cryoPROS refinement module. The quality of the generated particles inherently impacts subsequent operational efficiency and the feasibility of achieving credible results. Fig. 1 illustrates the impressive fidelity of cryoPROS-generated particles in mimicking the signal characteristics of authentic cryo-electron microscopy (cryo-EM) data. It's noteworthy that these generated particles exhibit a uniform distribution of orientations.

### 3 Importance of micelle reconstruction in cryoPROS-MP

We present the results derived from the application of cryoPROS to the membrane protein $Na_X$. For this study, the samples were suitably embedded in a micelle. The two-dimensional classification averages of particle images obtained through cryoPROS are given in Fig. 2a. Additionally, we provide the reconstructed density maps produced by both cryoPROS and cryoPROS-MP methods, displayed in Fig. 2b and Fig. 2c, respectively. A comparative analysis of these results underscores the enhanced performance offered by cryoPROS-MP. This superior performance firmly substantiates the incorporation of the additional "micelle reconstruction" module within this method. These promising findings not only validate the efficacy of cryoPROS-MP but also open a new avenue for its application in tackling more complex cases in the future.

### 4 Robustness analysis of cryoPROS and cryoPROS-MP

Besides the input raw particle images, both cryoPROS and cryoPROS-MP depend on an initial latent volume. Currently, this volume is derived by lowpass-filtering the density map of a homolog. Consequently, it's pivotal to examine the robustness of cryoPROS and cryoPROS-MP in relation to this initial latent volume. Specifically, we must assess whether performance is influenced by the choice of homologous protein and the lowpass filter's cutoff frequency. In the subsequent sections, we analyze the impact of different homologous proteins and various lowpass filter cutoff frequencies on the results.

#### 4.1 Effect of the choice of homologous protein.

Our experimental design involved the deployment of three distinct homologous proteins within both the HA trimer and $Na_X$ datasets. For the HA trimer, the abundance of readily available homologous proteins provided an intriguing opportunity for investigation. We selected homologous proteins with varying degrees of sequence identity—67%, 47%, and 17% relative to the target model, in order to thoroughly probe the impact of sequence divergence on cryoPROS performance. Remarkably, despite these differing sequence identities, the quantitative results of the reconstructed density maps in Table 2 exhibit comparable performance.

In contrast, for the $Na_X$ dataset, the lack of homologous proteins with various sequence similarities posed a major challenge. As a result, our methodology involved choosing homologous proteins with relatively closer sequence identities to the target model. Despite the limited range of homologous proteins, our results maintained similar model-to-map resolutions for the reconstructed density maps (see Table 2). The above two experiments establish the robustness of the proposed cryoPROS and the enhanced version cryoPROS-MP with respect to homologous proteins.

#### 4.2 Effect of the initial volume's lowpass filter cutoff frequency.

The latent volume in cryoPROS is initialized by the lowpass filtering homologous protein, and we test the robustness of cryoPROS by varying the cutoff frequency (Nyquist, 10Å, 20Å, 30Å) of the lowpass filter using two homologous proteins (PDB ID: 2RFU and 6IDD) in the HA Trimer dataset (EMPAIR-10096). The atomic model of the target protein solved from the data (EMPAIR-10097), via tilt-collection strategy is listed in PDB as 3WHE.

The sequence identity between the target (PDB ID: 3WHE) and the homolog (PDB ID: 2RFU) stands at 17%, while between the target (PDB ID: 3WHE) and another homolog (PDB ID: 6IDD), it's 47%.

Using homolog 2RFU as the initial map, the resulting map from CryoSPARC was lowpass filtered at different cutoff frequencies, including Nyquist, 10Å, 20Å, and 30Å. These are designated as cryoPROS-2RFU-Nyquist up to cryoPROS-2RFU-LP30. To assess the spectral similarity to the homologous protein, the cryoPROS-generated maps, along with the target map, were evaluated using the model (model 2RFU)model-to-map FSC, as depicted in Fig. 3a-i. With the exception of cryoPROS-2RFU-Nyquist, all other FSC curves intersect the 0.5 thresholds at frequencies below 25Å. This suggests that cryoPROS doesn't gather information from the initial volume at frequencies exceeding 25Å, if a 10Åor lower lowpass cutoff frequency is taken. Thus, we conclude that the model bias has not been introduced in cryoPROS.

To assess the spectral similarity to the target protein, the cryoPROS-generated maps were evaluated using the model (3WHE)-to-map FSC, as shown in Fig. 3a-ii. While the cutoff frequency of 20Åslightly underperforms compared to 10Å, the performance at 30Åis significantly worse. Based on these findings, we conclude that for this dataset, a lowpass filter cutoff frequency of 10Åis the most suitable choice.

Meanwhile, the homolog 6IDD has a sequence similarity of 47% with the target. Using 6IDD as the initial volume, parallel experiments reaffirm our findings (refer to Fig. 3iia-b), strengthening the reliability of our observations. Consistent with our earlier results, a lowpass filter cutoff frequency of 10Å effectively prevents information with a spectral frequency higher than 21Å from influencing the cryoPROS results. Interestingly, cryoPROS-6IDD-Nyquist does not resemble the target protein. This suggests that for a homolog with a high sequence similarity to the target, it is prudent to apply a lowpass filter before using it as the initial volume.

Undoubtedly, the choice of this parameter warrants exploration by users across different datasets. Our observations indicate that while various low-pass filter parameters in CryoPROS don't capture distinct characteristics of homologous proteins, there are discernible differences in analytical precision. However, when balancing the need for low model bias and high analytical accuracy, a 10Å lowpass filter for the cryoPROS initial latent volume emerges as optimal. As highlighted in the main text, a consistent 10Å-low pass setting was adopted for all datasets. The rationale for this choice has been validated in previous sections, where the reliability of CryoPROS results was confirmed under this specific configuration.

### 5 Model bias-related experiments

#### 5.1 Computation of KL divergence and SNR.

To demonstrate the superiority of cryoPROS-generated noisy particles over those generated by the Gaussian noise substitution method, we compute the KL divergence between the generated and real particles and the particle SNR. Let $\{\mathbf{x}_i^g\}_{i=1}^N$ and $\{\mathbf{x}_i^r\}_{i=1}^N$ represent the sets of generated and real particles, respectively, with $N$ denoting the number of particles. To compute the KL divergence, we first convert the particle images to probability histograms, denoting the number of bins as $B$, and for a particle image $\mathbf{x}$, its probability histogram $\mathbf{p} \in \mathbb{R}^B$. Let the probability histograms for $\{\mathbf{x}_i^g\}_{i=1}^N$ and $\{\mathbf{x}_i^r\}_{i=1}^N$ be represented as $\{\mathbf{p}_i^g\}_{i=1}^N$ and $\{\mathbf{p}_i^r\}_{i=1}^N$, respectively. Then, the average KL divergence for all particles is defined as:

$$\mathrm{KL}(\{\mathbf{x}_i^g\}_{i=1}^N, \{\mathbf{x}_i^r\}_{i=1}^N) = \frac{1}{N}\sum_{i=1}^{N}\sum_{j=1}^{B}\mathbf{p}_{ij}^g \log\frac{\mathbf{p}_{ij}^g}{\mathbf{p}_{ij}^r} \tag{1}$$

where $\mathbf{p}_{ij}^g$ and $\mathbf{p}_{ij}^r$ denotes the $j$-th component of $\mathbf{p}_i^g$ and $\mathbf{p}_i^r$, respectively. In practice, since the pixel values within a particle image remain unbounded, we first clip the pixel values within the range of $[-4, 4]$, and the number of bins $B$ is set to 1024.

We estimate the SNR of noisy particles using the method proposed in [1]. First, we randomly sample $N$ noisy particles, denoted as $\{\mathbf{x}_i\}_{i=1}^N$. Subsequently, we randomly sample $N$ background particles from the original micrographs, denoted as $\{\mathbf{x}_i^b\}_{i=1}^N$, which contain only pure noise. Denote the mean and variance for each background particle $\mathbf{x}_i^b$ as $\boldsymbol{\mu}_i^b$ and $\mathbf{v}_i^b$, respectively. Then, we normalize the noise in $\mathbf{x}_i$ and convert it to $\hat{\mathbf{x}}$ through the equation: $\hat{\mathbf{x}}_i = \mathbf{x}_i - \boldsymbol{\mu}_i^b$. We denote the mean and variance for $\hat{\mathbf{x}}_i$ as $\boldsymbol{\mu}_i$ and $\mathbf{v}_i$. The average SNR (dB) for the noisy particles is then defined as:

$$\mathrm{SNR} = \frac{10}{N} \sum_{i=1}^{N} \log_{10}(\mathbf{v}_i) - \log_{10}\left(\mathbf{v}_i^b\right).$$

In practice, we set the value of $N$ to 10.

#### 5.2 Gaussian noise substitution method.

In this section, we aim to establish the necessity of employing Conditional Variational Autoencoders (CVAE) by comparing it with an alternative approach: Gaussian noise substitution method. The Gaussian noise substitution method involves substituting the learned noise in cryoPROS with Gaussian noise while keeping all other parameters unchanged. The step-by-step process of this alternative method is as follows: (1) Initialization: The lowpass filtered homologous protein serves as the initial latent volume for projection, utilizing a uniform pose direction, where the projection process was carried out using the `relion_project` module within Relion. (2) Modulation and noise addition: CTF modulation is applied to the projected images, followed by the addition of Gaussian noise (standard deviation of added white Gaussian noise is 50, resulting in a signal-to-noise ratio (SNR) of −14.84dB). (3) Integration into refinement module: Gaussian noisy articles are imported into cryoPROS's refinement model, and the same processing steps are repeated. Subsequently, the latent volume is updated, initiating an iterative cycle of the aforementioned steps. We evaluate these two methods on HA-trimer and $\mathrm{Na_X}$ datasets, and the results are shown in Table 3. From the table, we find the Gaussian noise substitution method failed to achieve the desired quality on the $\mathrm{Na_X}$ dataset, and for the HA trimer dataset, the Gaussian noise substitution method exhibits some potential for success, whereas the reconstructed density displayed a closer resemblance to the homologous protein than the result produced by cryoPROS, which raises concerns about the potential model bias problem.

In summary, from the results of this series of experiments, the Gaussian noise substitution method proves to be inadequate for mitigating model bias and generating accurate protein reconstructions and shows the indispensable role played by the cryoPROS generative module in averting model bias effectively and producing reliable outcomes in cryo-EM studies.

Our investigation involves conducting experiments on two distinct datasets, namely the HA-trimer and $Na_X$. The obtained results unveil intriguing insights. The Gaussian noise substitution method does not yield satisfactory outcomes for the $Na_X$ protein (see Table 3). The resulting reconstructions fail to achieve the desired quality, indicating the ineffectiveness of the Gaussian noise substitution method in this context. On the other hand, in the case of the HA trimer, the Gaussian noise substitution method exhibits some potential for success. However, closer examination reveals that the outcomes bear a striking resemblance to the homologous protein, rather than the results produced by cryoPROS (see Table 3). This phenomenon indicates a notable risk of introducing model bias into the reconstructions through the Gaussian noise substitution method.

In summary, these series of experiments convincingly underscore the significance of the generative module within the cryoPROS framework. The Gaussian noise substitution method proves to be inadequate for mitigating model bias and generating accurate protein reconstructions. The results emphasize the indispensable role played by the cryoPROS generative module in averting model bias effectively and producing reliable outcomes in cryo-electron microscopy studies.

##### 5.3 Model bias verification of other datasets.

In main text, we employed a structural refinement method to validate the accuracy of cryoPROS resulting map in the case of HA trimer. This validation demonstrated a remarkable closeness between the cryoPROS outcomes and the true structure, while simultaneously showcasing a significant dissimilarity from homologous proteins structure. This evidence substantiates the claim that cryoPROS is devoid of model bias risks.

Expanding upon this notion, we extended our investigation to encompass three additional datasets: SIM2 (see Fig. 4a), the PO-subset of EM-10024 (see Fig. 4b), and $Na_X$ (see Fig. 4c). We found that the experiment had consistent results across different data sets. These results further reinforce our earlier assertion about cryoPROS's proficiency in generating unbiased results. It reinforces its capability to produce accurate reconstructions that remain faithful to the target structures, while mitigating the risks associated with model bias.

#### 6 Hierarchical structure of CVAE

In cryoPROS, a hierarchical structure [2, 3] is adopted for the CVAE consisting consists of $L$ stochastic layers. Specifically, we represent the latent variable as

$\mathbf{z} = (\mathbf{z}^1, \ldots, \mathbf{z}^L)$, and the prior model is decomposed as follows:

$$p(\mathbf{z}|\boldsymbol{\Theta}) = p(\mathbf{z}^L|\boldsymbol{\Theta}) \prod_{l=1}^{L-1} p(\mathbf{z}^l|\mathbf{z}^{>l}, \boldsymbol{\Theta}) \tag{2}$$

where $\mathbf{z}^{>l} = (\mathbf{z}^{l+1}, \ldots, \mathbf{z}^L)$. Similarly, the inference models follow the same order:

$$q(\mathbf{z}|\mathbf{x}, \boldsymbol{\Theta}) = q(\mathbf{z}^L|\mathbf{x}, \boldsymbol{\Theta}) \prod_{l=1}^{L-1} q(\mathbf{z}^l|\mathbf{z}^{>l}, \mathbf{x}, \boldsymbol{\Theta}). \tag{3}$$

Then, the cELBO can be expressed as:

$$\begin{aligned} \text{cELBO} =&\mathbb{E}_{q(\mathbf{z}|\mathbf{x},\boldsymbol{\Theta})} \log p(\mathbf{x}|\mathbf{z}, \boldsymbol{\Theta}) - \text{KL}(q(\mathbf{z}^L|\mathbf{x}, \boldsymbol{\Theta}) \| p(\mathbf{z}^L|\boldsymbol{\Theta})) \\ &- \sum_{l=1}^{L-1} \mathbb{E}_{q(\mathbf{z}^{>l}|\mathbf{x},\boldsymbol{\Theta})} \text{KL}(q(\mathbf{z}^l|\mathbf{z}^{>l}, \mathbf{x}, \boldsymbol{\Theta}) \| p(\mathbf{z}^l|\mathbf{z}^{>l}, \boldsymbol{\Theta})). \end{aligned} \tag{4}$$

Regarding the inference model $q(\mathbf{z}|\mathbf{x}, \boldsymbol{\Theta})$, we assume the following form:

$$q(\mathbf{z}^l|\mathbf{z}^{>l}, \mathbf{x}, \boldsymbol{\Theta}) = \mathcal{N}(\mu_q^l(\mathbf{a}^l, \mathbf{b}^l, h_\theta^l(\mathbf{v})), \sigma_q^l(\mathbf{a}^l, \mathbf{b}^l, h_\theta^l(\mathbf{v}))), \quad l = 1, 2, \ldots, L \tag{5}$$

where $\mathbf{a}^l$ and $\mathbf{b}^l$ denotes the encoding and decoding feature in $l$-th layer, respectively. $\mathbf{v} = \mathbf{C}(\psi)P(\phi)\mathbf{V}_{latent}$ denotes the projection of the latent volume, and it is embedded into the latent space using a neural network denoted as $h_\theta(\cdot)$. $\mu_q^l$ and $\sigma_q^l$ are networks that convert $(\mathbf{a}^l, \mathbf{b}^l, h_\theta^l(\mathbf{v}))$ to the parameters of a Gaussian distribution. The encoding features $\{\mathbf{a}^l\}_{l=1}^L$ are recursively obtained as follows:

$$\mathbf{a}^1 = f_\theta^1(\mathbf{x}), \quad \mathbf{a}^l = f_\theta^l(\mathbf{a}^{l-1}), \quad l = 2, \ldots, L, \tag{6}$$

where $f_\theta^l$ represents the convolutional block in the $l$-th encoding layer. The decoding features $\mathbf{b}^l$ are obtained through the recursion:

$$\mathbf{b}^{l-1} = g_\theta^l(\mathbf{z}^l, \mathbf{b}^l), \quad l = 2, \ldots, L, \tag{7}$$

where $\mathbf{z}^l$ is sampled from $\mathcal{N}(\mu_q^l(\mathbf{a}^l, \mathbf{b}^l, h_\theta^l(\mathbf{v})), \sigma_q^l(\mathbf{a}^l, \mathbf{b}^l), h_\theta^l(\mathbf{v}))$, $\mathbf{b}^L$ is a constant vector that is set as a learnable parameter, and $g_\theta^l$ is the convolutional block in $l$-th decoding layer. Additionally, for the prior model $p(\mathbf{z}|\boldsymbol{\Theta})$, we assume the following form:

$$p(\mathbf{z}^l|\mathbf{z}^{>l}, \boldsymbol{\Theta}) = \mathcal{N}(\mu_p^l(\mathbf{b}^l, h_\theta^l(\mathbf{v})), \sigma_p^l(\mathbf{b}^l, h_\theta^l(\mathbf{v}))), \quad l = 1, 2, \ldots, L, \tag{8}$$

and the generative model $p(\mathbf{x}|\mathbf{z}, \boldsymbol{\Theta})$ is assumed to be $\mathcal{N}(g_\theta^1(\mathbf{z}^1, \mathbf{b}^1), \mathbf{I})$. We adopt the Residual Dense Block (RDB) [4] as our convolutional block for $f_\theta^l$ and $g_\theta^l$, and the model architecture is shown in Fig. 5. The loss function for the CVAE aims to minimize the −cELBO, and with the given parametrization, the loss function is:

$$\mathrm{Loss}(\mathbf{x}, \mathbf{\Theta}) = \mathbb{E}_{q(\mathbf{z}|\mathbf{x},\mathbf{\Theta})} \|g^1_\theta(\mathbf{z}^1, \mathbf{b}^1) - \mathbf{x}\|_2^2 + \sum_{l=1}^{L} \mathrm{KL}\left(\mathcal{N}(\mu^l_q, \sigma^l_q) \| \mathcal{N}(\mu^l_p, \sigma^l_p)\right). \quad (9)$$

The KL divergence for two Gaussian distributions has an analytical form, which is:

$$\begin{aligned}\mathrm{KL}\left(\mathcal{N}(\mu_q, \sigma_q) \| \mathcal{N}(\mu_p, \sigma_p)\right) = &\frac{1}{2}\left(\log(|\sigma_p|/|\sigma_q|) - K\right. \\ &\left.+ \mathrm{Tr}(\sigma_p^{-1}\sigma_q) + (\mu_q - \mu_p)^T \sigma_p^{-1}(\mu_q - \mu_p)\right),\end{aligned} \quad (10)$$

where $K$ denotes the dimension of the latent variable $\mathbf{z}$.

| Datasets | SIM1 | SIM2 | EM-10024 | PO-subset of EM-10024 | HA Trimer | $Na_X$ |
|---|---|---|---|---|---|---|
| Data Source | Simulation | Simulation | EMPIAR repository | Manual selection | EMPIAR repository | Collected in-house |
| Particles | 130,000 | 130,000 | 43,585 | 14,436 | 130,000 | 411,520 |
| Preferred orientation | No | Yes | Yes | Yes | Yes | Yes |
| Target Protein | HA in H3N2 influenza A viruses | HA in H3N2 influenza A viruses | TRPA1 ion channel | TRPA1 ion channel | HA in H3N2 influenza A viruses | $Na_X$ |
| PDB ID (target protein) | 3WHE | 3WHE | 3J9P | 3J9P | 3WHE | None |
| Latent volume source for cryoPROS iter1 | None | Homolog Protein | None | Alphafold2 predicted | Homolog Protein | Homolog Protein |
| Latent volume for cryoPROS iter1 | None | HA in Influenza B virus | None | Rat TRPA1 | HA in H7N9 influenza A viruses | Nav1.6 |
| Resolution obtein-ed by cryoPROS | None | 3.28Å | None | 8.4Å | 3.90Å | 7.22Å |

**Table 1**: **Attributes of simulated and authentic cryo-EM datasets appeared in the main text**

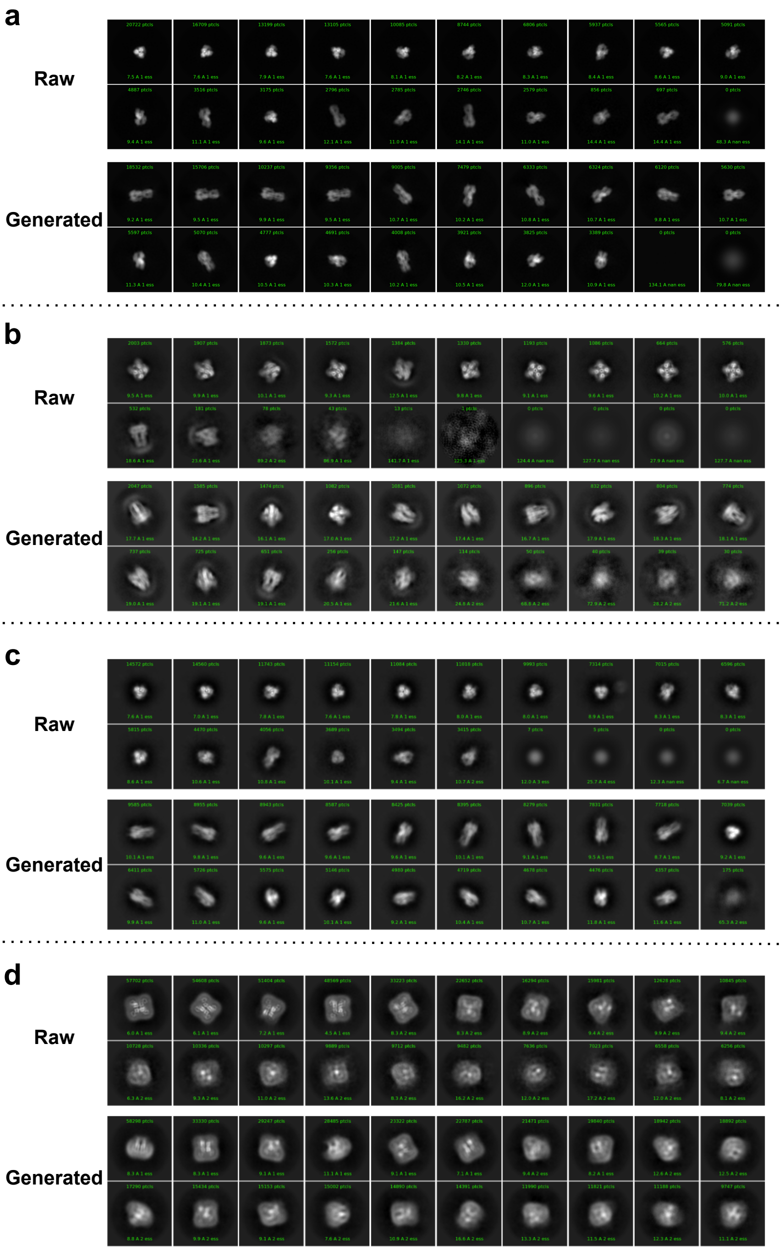


**Fig. 1**: **Comparative analysis of raw and generated particles.** Averaged 2D classification comparison illustrating the similar signal and different pose distributions between raw particle data (upper row) and generated particles (lower row) across preferred oriented datasets, panels a to b, including SIM2, PO-subset of EM-10024, HA trimer, and $Na_X$.

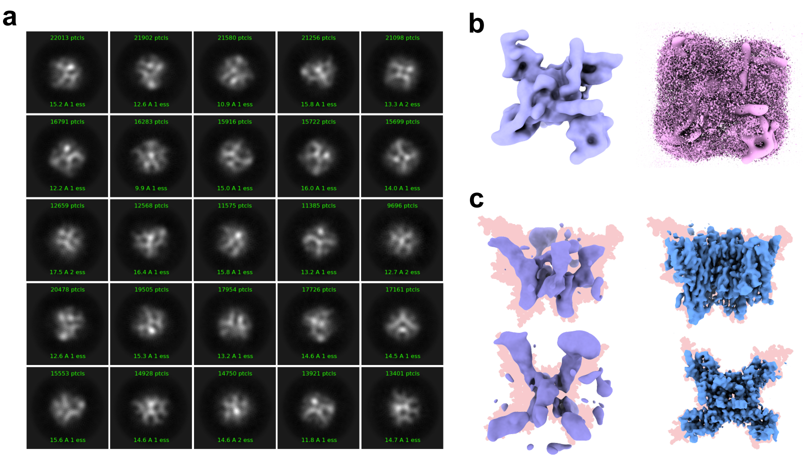


**Fig. 2**: **Necessity of Micelle Reconstruction Step in CryoPROS-MP.** **a**, Two-dimensional classification average obtained by excluding the micelle reconstruction step in CryoPROS-MP. The comparison illustrates the effects of this omission on particle generative quality. **b**, Visualization of latent volumes before and after the micelle reconstruction step. **c**, Resulting map comparison between CryoPROS (left) and CryoPROS-MP (right), highlighting the improvements achieved by the "micelle reconstruction" module in enhancing structural accuracy and detail.

| Dataset | PDB ID (sequence similarity) | Half-map resolution (Å) | Model-to-map resolution (Å) | Average Q-score |
|---|---|---|---|---|
| HA trimer | 5XL8 (67%) | 3.35 | 4.47 | 0.395 |
| | 6IDD (47%) | 3.27 | 3.90 | 0.435 |
| | 2RFU (17%) | 4.35 | 3.35 | 0.388 |
| Na$_X$ | 7XM9 (58%) | 3.37 | 7.22 | 0.301 |
| | 8FHD (56%) | 3.33 | 7.22 | 0.325 |
| | 6AGF (56%) | 3.34 | 7.22 | 0.321 |

**Table 2**: **The performance of cryoPROS using different initial homologous proteins.**

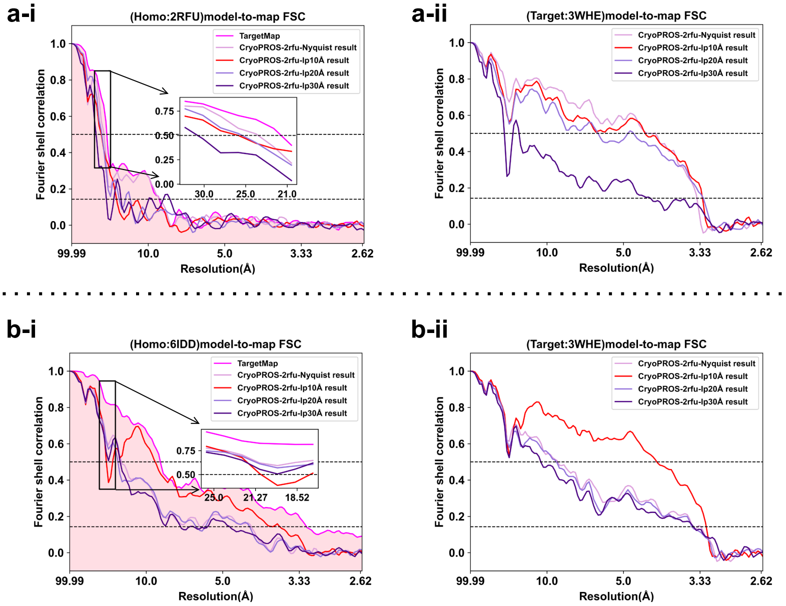


**Fig. 3**: **Evaluating CryoPROS performance using different lowpass filter cutoff frequencies for the initial volume. a-i**, When employing homolog 2RFU as the initial map, we lowpass filtered the resulting map from CryoSPARC using various cutoff frequencies: Nyquist, 10Å, 20Å, and 30Å. These configurations are labeled from cryoPROS-2RFU-Nyquist to cryoPROS-2RFU-LP30. For spectral similarity assessment relative to the homologous protein, we analyzed the cryoPROS-derived maps, in conjunction with the target map, through the model-to-map FSC based on the 2RFU model. The region close to the 0.5 threshold of the FSC curves is magnified and emphasized within the box. **b-i**, Analogous to **a-i**, this illustrates the spectral similarity assessment with respect to the target protein (3WHE) rather than the homologous protein (2RFU). **b**, **b** mirrors **a** (covering both **i** and **ii**), with the distinction being that 6IDD serves as the initial model in place of 2RFU.

| Method | HA trimer | | $Na_X$ | |
|---|---|---|---|---|
| | Target model (3WHE) | Homolog model (6IDD) | Target model ($Na_X$) | Homolog model (Nav1.6: 8FHD) |
| GN substitution | 4.14 | 9.58 | 40.23 | 31.28 |
| CryoPROS | 4.14 | 20.96 | 7.22 | 23.47 |

**Table 3**: **Comparison of model-to-map resolutions (Å) between density maps reconstructed using Gaussian noise substitution method and cryoPROS. Higher resolution is preferred for the target model, while lower resolution is preferred for the homolog model.**

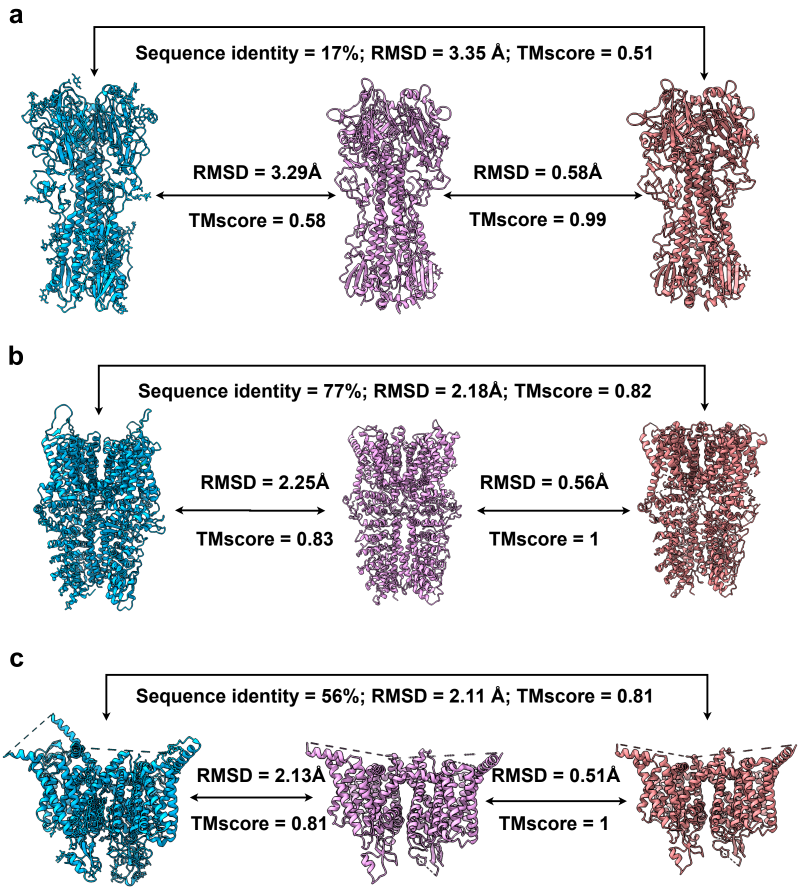


**Fig. 4**: **Structure refinement experiments on other datasets. a**, For the SIM2 dataset, a comparative analysis is presented involving the atomic models of the homologous protein, a structure-refined model derived from the cryoPROS resulting map (based on target model), and the target protein. The similarity between the two models is evaluated through Root Mean Square Deviation (RMSD) and TMscore calculations. **b,c**, Similar results are observed for the PO-subset of EM-10024 and $Na_X$ datasets.

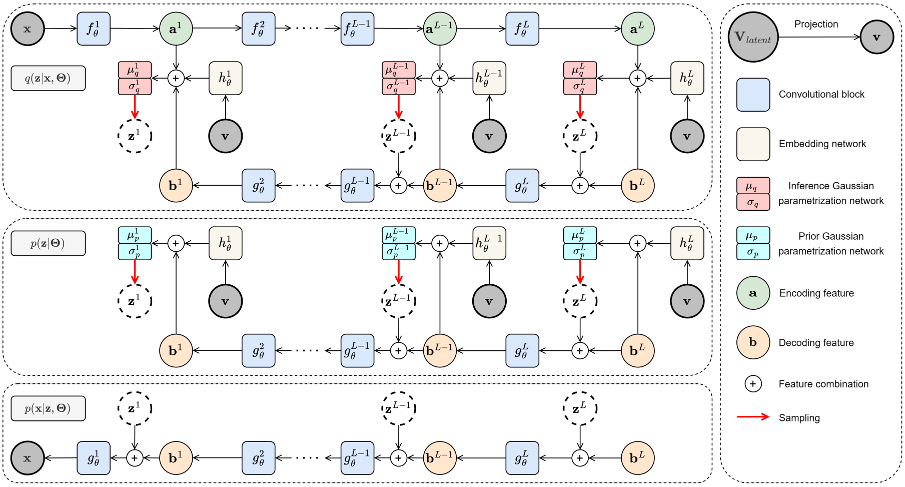


**Fig. 5**: **Network architecture of the hierarchical CVAE model, including the inference model $q(\mathbf{z}|\mathbf{x}, \boldsymbol{\Theta})$, the prior model $p(\mathbf{x}|\boldsymbol{\Theta})$, and the generative model $p(\mathbf{x}|\mathbf{z}, \boldsymbol{\Theta})$.**
